# Optimizing Post-Transplantation Detection of Subcutaneously Transplanted Islets Using Dithizone Staining

**DOI:** 10.64898/2026.01.06.697969

**Authors:** James Lu, Matthew Ishahak, Marlie M. Maestas, Jeffrey R. Millman

**Affiliations:** Division of Endocrinology, Metabolism and Lipid Research, Washington University School of Medicine, St. Louis, MO, USA

**Keywords:** Islet Transplantation, Diabetes, SC-islets, Dithizone Staining

## Abstract

**Introduction:** Pancreatic islet transplantation is a promising therapeutic strategy to restore insulin independence in patients with type 1 diabetes mellitus. In addition to primary human islets, human pluripotent stem cell-derived islets have shown clinical promise. However, optimizing transplantation sites and improving methods for standardized islet dosing and post-transplant graft localization are key limitations. The subcutaneous space offers an alternative site for transplantation due to its accessibility and minimally invasive nature; nevertheless, poor vascularization and difficulty locating engrafted islets limit its experimental utility. The use of dithizone, a zinc-chelating dye that selectively binds the high intracellular zinc concentrations of insulin-producing β-cells, may enable rapid graft localization through selective staining. Here, we investigate whether dithizone staining can reliably localize transplanted islet grafts within the subcutaneous space of murine models.

**Methods:** SC-islets were generated from HUES8 stem cells and functionally validated using glucose-stimulated insulin secretion. Islet volume was standardized prior to transplantation using automated islet equivalent quantification with the BioRep Islet Cell Counter. Approximately 4,000 islet equivalents of either SC-islets or primary human islets were transplanted into the subcutaneous space of immunodeficient mice. Graft function was assessed longitudinally via blood glucose monitoring, glucose tolerance testing, and human C-peptide measurements. Four weeks post-transplantation, dithizone staining was applied to harvested skin tissue to localize islet grafts, and immunohistochemical validation was performed.

**Results:** Automated quantification of islet equivalents enabled consistent dosing across transplantation groups. Transplanted islets maintained function *in vivo*, as demonstrated by reduced blood glucose levels, preserved glucose tolerance, and detectable human C-peptide. Dithizone staining produced clear and selective red labeling of insulin-producing β-cell grafts within the subcutaneous tissue, enabling reliable localization and recovery of engrafted islets. Immunohistochemical analyses confirmed the presence of insulin-expressing cells at isolated islet grafts.

**Discussion:** Dithizone staining is a rapid, cost-effective approach to identify subcutaneous islet grafts and enables downstream analyses, addressing a significant limitation in islet transplantation research. When combined with standardized pre-transplant islet quantification, this approach provides a cohesive and reproducible framework for evaluating subcutaneous islet transplantation. Future studies could complement these results by assessing the potential effects of dithizone exposure and exploring its applicability across other transplantation sites and imaging modalities.

## INTRODUCTION

Insulin-secreting pancreatic islet transplantation has emerged as a promising strategy for restoring insulin independence in patients with type 1 diabetes (T1D) mellitus (1). For over two decades, transplantation protocols have largely focused on the infusion of primary human islets, isolated from the pancreas of deceased donors, via the hepatic portal vein, a site that offers immediate access to the systemic circulation and mimics endogenous insulin delivery to the liver (2–4). More recently, human pluripotent stem cell-derived islets (SC-islets) have been developed and shown promise not only in preclinical animal transplantation models (5–9) but also in a clinical trial (10). However, despite early clinical success, SC-islet transplantation is still limited by several drawbacks, including poor long-term graft survival, blood-mediated inflammatory responses, immune-mediated graft destruction, and challenges in monitoring graft viability and function over time (11). Furthermore, the invasive nature of hepatic portal vein transplantations complicates both the retrieval of islets and their longitudinal assessment, which hinders mechanistic studies and optimization of graft survival strategies (12).

To overcome these limitations, preclinical studies have been exploring alternative transplantation sites in murine models that offer improved accessibility, physiological support, and experimental flexibility. Among these, the renal subcapsular space (kidney capsule), intramuscular region, and subcutaneous cavity have gained particular interest (13). The kidney capsule remains a popular site due to its high vascularization and ease of access in small animal models but has limited translational capacity and is not a suitable location for human transplantations, which constrains its utility (14). The intramuscular space allows for better vascular integration but may involve complex surgical implantation and fibrotic encapsulation (15). The subcutaneous space, by contrast, represents an especially attractive transplantation site due to minimally invasive access, potential for real-time monitoring, non-terminal graft retrieval, and device compatibility (16,17). However, this space is relatively poorly vascularized, which presents a challenge to islet engraftment and survival (18). Furthermore, reliably locating and assessing transplanted islets in the subcutaneous space is very difficult due to graft dispersion and limited visual contrast with surrounding tissue (19).

In addition to challenges related to graft localization, a major technical limitation in both preclinical and translational islet transplantation studies is the accurate and reproducible quantification of islets prior to transplantation. Standardization of transplanted islet volume or islet equivalents (IEQs) is essential for interpreting graft survival, functional outcomes, and dosage responses across experimental groups (20,21). However, manual IEQ quantification is labor-intensive and impractical when processing large numbers of samples. These limitations can introduce significant variability and reduce experimental reproducibility (22–24). The BioRep Islet Cell Counter (ICC), an automated image-based islet counter, offers a standardized and reproducible approach to IEQ quantification by objectively measuring islet size distribution and total islet volume (25). Despite its growing adoption in islet research, integration of the BioRep ICC with transplantation and dosage response relationships remains underreported, particularly in the context of subcutaneous transplantation models.

Beyond the need for standardized islet quantification, reliable post-transplant identification of grafted islets remains a significant challenge, particularly within poorly vascularized and spatially diffuse sites such as the subcutaneous space. Dithizone (DTZ), a zinc-chelating dye that binds selectively to the high intracellular zinc concentrations found in insulin-producing β-cells, has been widely used *in vitro* to assess islet purity and viability (26,27). The staining has a distinct red color that provides a convenient and specific marker for visually identifying pancreatic islets. Despite its frequent application in islet isolation protocols, the application of DTZ for *in vivo* identification of engrafted islets has not been well explored. Notably, the potential of DTZ as a non-destructive tool for locating islets within the subcutaneous space after transplantation has not been fully investigated.

In this study, we investigate the application of DTZ staining as a method for detecting and localizing transplanted islet grafts within the subcutaneous space of murine models. We hypothesize that DTZ can serve as a non-destructive, rapid, and cost-effective method for localizing transplanted islets, thereby addressing a key limitation in the post-transplantation assessment of the subcutaneous site. When combined with standardized pre-transplant islet quantification using automated islet equivalent measurements, we aim to provide a methodological advancement that supports preclinical islet transplantation studies.

## MATERIALS AND METHODS

### Ethics Statement

This work has been approved by the Washington University School of Medicine, including Institutional Biological & Chemical Safety Committee (20886), Stem Cell Research Oversight Committee (17–005), and Institutional Animal Care and Use Committee (IACUC; Protocol #24-0185).

### SC-islet Differentiation

HUES8 stem cell line was generously provided by Dr. Douglas Melton (Harvard University). SC-islets were differentiation from stem cells following our previously published protocol (28). Briefly, undifferentiated stem cells were seeded onto a Matrigel (Corning; 354277) treated tissue culture flask at 0.63 × 10^6^ cells/cm^2^ with mTesR1 (StemCell Technologies; 05850) and 10 μM Y-27632 (Pepro Tech; 129382310MG). After 24 h, media and growth factors were added to start the differentiation process (Supplementary Table 1). On Stage 6 Day 7, the differentiation was dispersed with TrypLE at 0.2 ml cm^−2^ (Gibco; 12-604-013) and seeded into individual wells of a 6-well dish with 4-mL of ESFM media on an Orbi-Shaker (Benchmark) at 100 RPM. After 6-12 more days, cells were used for assessment. All factors, timing, and media formulations for the differentiation are in Supplementary Table 1. Light Microscopy. Light Microscopy images were taken using an inverted light microscope (Leica DMi1).

### Primary Human Islet Culture

Primary human islets were purchased from Prodo Labs. Upon arrival, islets were centrifuged for 3 minutes at 300xg, then resuspended in CMRL (Mediatech; 99-603-CV) supplemented with 10% FBS (GE Healthcare; SH3007003). Primary islets were then cultured in individual wells of a 6-well dish with 5mL of CMRL+FBS media on an Orbi-Shaker (Benchmark) at 100 RPM for 2 days prior to transplantation. Donor islets came from a 55-year-old, white male with a BMI of 29.0.

### Glucose Stimulated Insulin Secretion Assay

SC-islets were washed with Krebs-Ringer bicarbonate buffer (KRB) composed of 128 mM NaCl, 5 mM KCl, 2.7 mM CaCl_2_ 1.2 mM MgSO_4_, 1 mM Na_2_HPO_4_, 1.2 mM KH_2_PO_4_, 5 mM NaHCO_3_, 10 mM HEPES (Gibco; 15630-080), and 0.1% bovine serum albumin. SC-islets were then transferred to transwells and equilibrated in KRB containing 2 mM glucose for 1 h at 37 °C. Following equilibration, the solution was replaced with fresh 2 mM glucose KRB for an additional 1 h, after which supernatant was collected to assess basal insulin secretion. The solution was subsequently replaced with 20 mM glucose KRB and incubated for a final 1 h before supernatant was collected. After the secretion assay, islets were dispersed into single cells using TrypLE and incubated for 30 minutes. Cells were counted using a Vi-Cell XR to normalize insulin secretion values to cell number. Insulin concentrations in collected supernatants were quantified using a human insulin ELISA (ALPCO; 80-INSHU-E10.1). All incubations were performed in a humidified incubator at 37°C with 5% CO_2_.

### Quantification of Islet Equivalents

Islet volume was standardized by quantifying islet equivalents using the BioRep ICC (BioRep Technologies; ICC4-115/230). Depending on the total islet yield, multiple counts were performed per well to ensure accurate quantification. For each count, a small aliquot of islets was transferred from the culture vessel into a BioRep ICC dish (BioRep Technologies; ICC-D2). A minimum of 200 µL of culture medium or phosphate-buffered saline was added to the dish to prevent desiccation and to allow islet clusters to remain suspended. The ICC dish was gently swirled to distribute and congregate islet clusters while avoiding significant overlap. Overlapping islets were minimized, as the BioRep ICC system identifies overlapping clusters as a single islet, which can lead to inaccurate IEQ measurements. Once the islets were adequately spaced, the ICC dish was placed into the BioRep counter. The “unstained” profile was selected, and red pixel calibration was manually overridden, as the islets were not labeled with any fluorescent stains. The system captured an image of the dish and generated a preview highlighting detected islet clusters. Non-islet debris was manually excluded prior to analysis. Upon confirmation, the software generated a report detailing total IEQs and islet size distribution.

### Subcutaneous Islet Transplantation

All animal studies were conducted in accordance with the guidelines and regulations approved by the Washington University in St. Louis’s Animal Care and Use Committee. Male NOD.Cg-*Prkdc^scid^*Il2rg*^tm1Wjl^/*SzJ mice were obtained from The Jackson Laboratory (005557) and randomly assigned to experimental groups (n = 5 per group). Mice were anesthetized prior to transplantation and injected with approximately 4,000 IEQs per mice. Islets were transplanted into the subcutaneous space on the dorsal medial left side using a 27-gauge needle (BD, 309623). Blood glucose levels were monitored using a Contour NEXT blood glucose Monitor (Bayer).

### Glucose Tolerance Test

For the glucose tolerance test (GTT), mice were fasted for 5-6 h prior to the assay and administered an intraperitoneal injection of glucose (3 g/kg body weight) dissolved in 0.9% sterile saline (Moltox, 51-405022.052). Blood glucose levels were measured at baseline and at 30-min intervals for a total duration of 120 min using a glucometer. C-peptide levels were measured using an ultrasensitive human C-peptide ELISA (Mercodia, 10-1141-01).

### Dithizone Staining

Several weeks after transplantation, mice were euthanized and the skin containing the subcutaneous graft site was harvested. Skin samples were placed in Petri dishes and submerged in a 1 mg/mL DTZ solution sufficient to fully cover the tissue. DTZ solution was prepared by first dissolving DTZ in dimethyl sulfoxide (Fisher Scientific; BP231-100), followed by dilution in phosphate-buffered saline and sterile filtration. Samples were incubated for 5 min at room temperature, after which DTZ-positive regions containing islet engraftments were identified visually, imaged with iPhone 15 Pro Max, and excised. Excised tissues were fixed in 4% paraformaldehyde for 24 h, transferred to 70% ethanol, and submitted to Histowiz for histological processing, including paraffin-embedding and sectioning onto microscope slides.

### Immunohistochemistry

Paraffin was removed using Histoclear (Electron Microscopy Sciences; 64111-04) followed by graded ethanol washes (Decon; 2716) to rehydrate the sections. Antigen retrieval was performed using a pressure cooker (Proteogenix; 2100 Retriever) in 0.05M EDTA buffer (Ambion; AM9261). Sections were blocked and incubated with primary antibodies diluted in immunostaining buffer consisting of 5% Donkey Serum (Jackson Immunoresearch; 017-000-121) and 0.1% Triton-X (Acros Organics; 327371000) in phosphate-buffered saline (Corning; 21-040-CV) overnight at 4°C in a humidified box. After washing, sections were incubated with appropriate secondary antibodies diluted in immunostaining buffer for 2 h at room temperature. Slides were mounted using DAPI Fluoromount-G (SouthernBiotech; 0100-20) and imaged on a Zeiss LSM 880 II Airyscan FAST confocal microscope. Image processing and analysis were performed using ImageJ.

## RESULTS

### Generation and Functional Validation of SC-islets

SC-islets are generated through a multistep differentiation protocol that recapitulates key stages of embryonic pancreatic development using temporally controlled growth factors and small molecules (Fig. 1A). To validate the proper differentiation of the islets prior to transplantation, we documented the morphology of the islets during the start of each key stage (Fig. 1B). To validate and ensure that islets were properly functioning, we performed a glucose-stimulated-insulin-secretion assay (GSIS) to confirm if the islets would be up to standard quality. Both primary human islets (4.144±0.055 µIU/10^3^ cells) and SC-islets (1.086±0.204 µIU/10^3^ cells) secreted insulin in response to high glucose (Fig. 1C). However, consistent with previous reports, SC-islets are functionally immature compared to primary (11). We confirmed that the cultured HUES8 SC-islets were up to standard quality and within the range of expected secretion levels in comparison to the primary human islets.

**Fig. 1.**
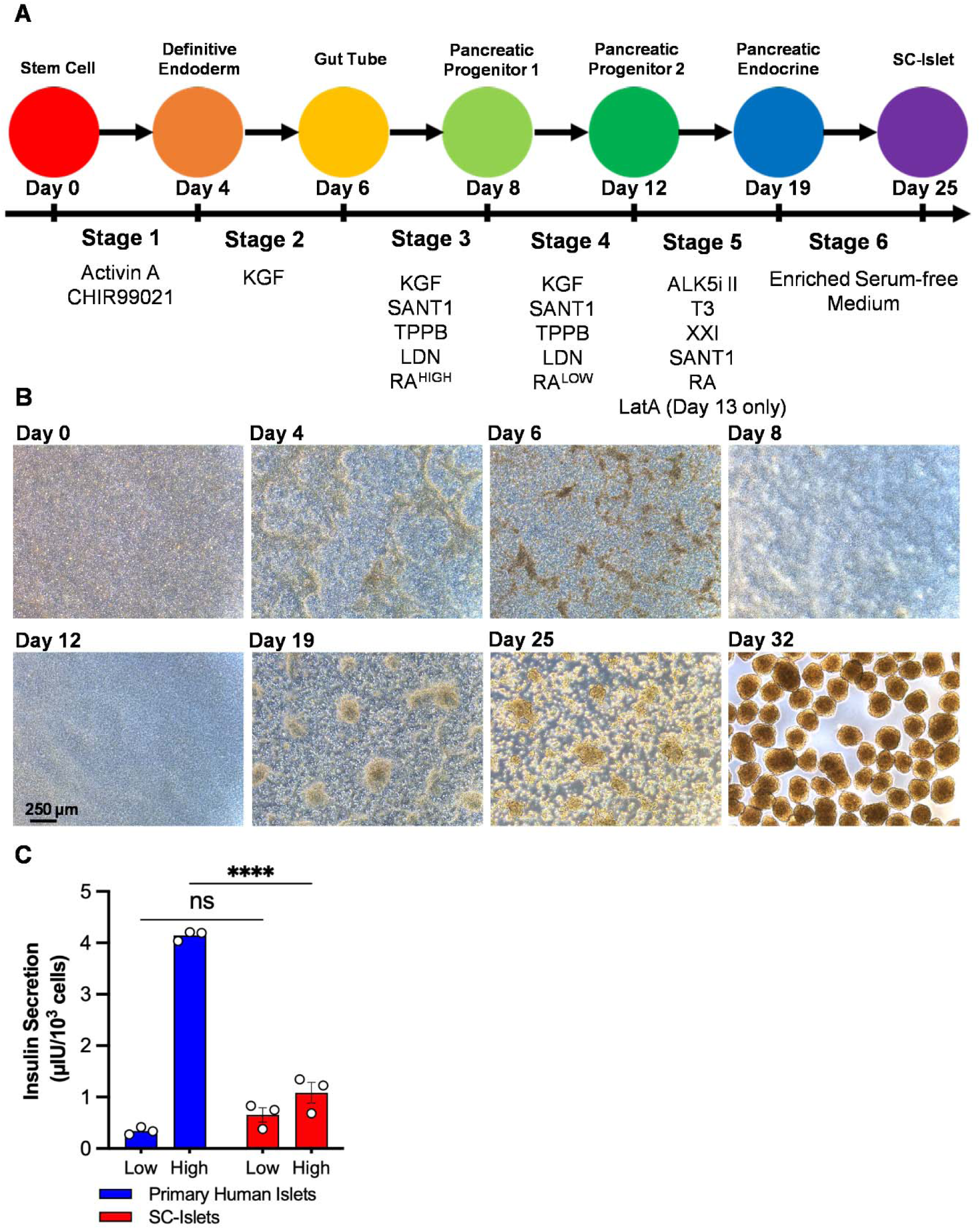
Generation and Functional Validation of SC-islets. (**A**) Schematic of stem cell differentiation protocol for making SC-islets. **(B)** Brightfield image highlighting morphological changes during stem cell differentiation into SC-islets. **(C)** Glucose stimulated insulin secretion test comparing the insulin sensitivity between primary human islets and SC-islets.

### Dosage Quantification and Islet Transplantation

We then measured the dosage of each transplantation with the ICC BioRep machine and made sure that the average amount of islets (primary human islets, 4200±227.7 IEQ; SC-islets, 4128±158.7 IEQ) transplanted was approximately the same (Fig. 2A-C). To further assess the quality of the islets and monitor their engraftment in the mice, we conducted weekly weight measurements and C-peptide assays (Fig. 2D-E). We observed no statistically significant differences in weight following transplantation, except at 4 weeks mice that received SC-islets were significantly lighter than non-transplanted mice (p = 0.004; Fig. 2D). We observed higher levels of C-peptide at 4 weeks post-transplantation in primary human islets (170.208±142.986 pmol/L) compared to SC-islets (59.247±15.433 pmol/L), but the differences were not significantly different (Fig. 2E). Similarly, we conducted mid-point and terminal GTT. There was no significant difference in the area under the curve for blood glucose levels following glucose injection at either 2-weeks or 4-weeks (Fig. 2F). These results demonstrated that the islets were properly functioning within the mice.

**Fig. 2.**
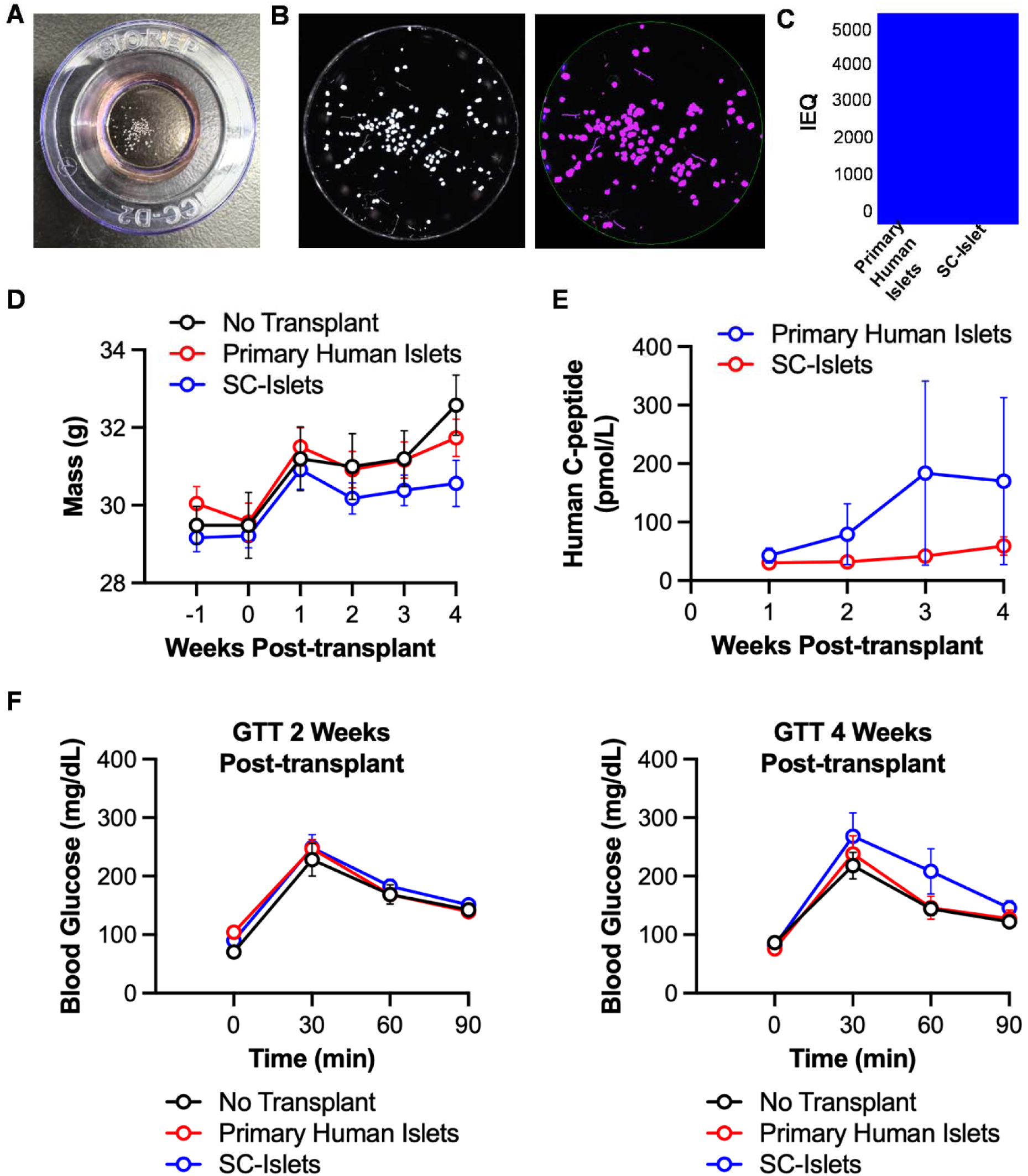
Dosage Quantification and Islet Transplantation. **(A)** Representative images of islets in the BioRep ICC dish before automated quantification of IEQs. **(B)** Representative images of binarized image of islets (left) and automatically detected islet clusters (right) by the BioRep ICC system and software. **(C)** IEQ comparison of the average dosage for each mice. **(D)** Weight of the mice during one before and up to 4 weeks after transplantion. **(E)** Comparison of the blood C-peptide levels between the primary human islets and SC-islets following transplation. **(F)** Glucose tolerance tests 2-weeks post-transplantation (left) and 4-weeks post-transplantation (right).

### Recovery and Identification of Islet Grafts

Four weeks after transplantation, mice were sacrificed and islet grafts were retrieved for assessment. To identify grafts within the subcutaneous space, we developed a methodology to stain transplanted islets using DTZ (Fig. 3A). Prior to DTZ staining, neither primary human islet nor SC-islet grafts were readily identifiable to the naked eye, making retrieval impractical (Fig. 3B). After soaking in a solution of 1 mg/mL DTZ for 5 minutes, grafts became visible to the eye due to the characteristic red color-staining of the zinc present in high amounts in β cells (Fig. 3C). Identified grafts were excised and immunohistochemistry (IHC) staining was performed to confirm that DTZ staining does not interfere with downstream assessments and to demonstrate its utility.

**Fig. 3.**
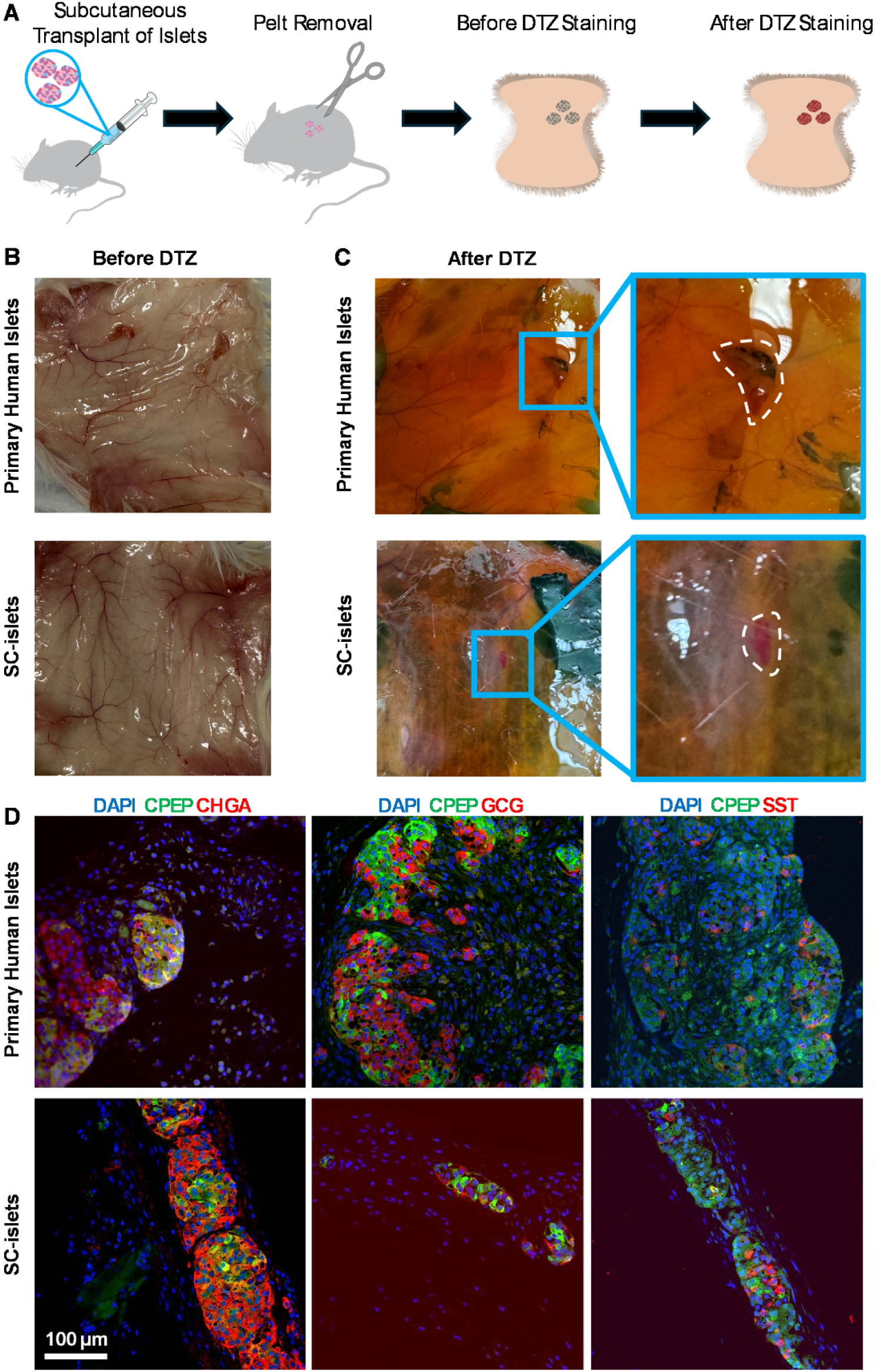
Recovery and Identification of Islet Grafts. (**A**) Schematic representation of the DTZ staining methodology. Cartoons retrieved from: NIAID Visual & Medical Arts. Syringe (bioart.niaid.nih.gov/bioart/505), Wood Mouse Silhouett (bioart.niaid.nih.gov/bioart/20), and Scissors (bioart.niaid.nih.gov/bioart/488) **(B-C)** Representative picture of the transplant location before (B) and after (C) DTZ staining highlighting engraftment of both human islets and SC-islets. **(D)** Immunofluorescent characterization of explanted primary human islet and SC-islet grafts following subcutaneous transplantation and DTZ staining.

Following IHC staining, both primary and SC-islets were positive of expected markers. Sections were stained with DAPI to label nuclei, C-peptide to identify insulin-producing β-cells, chromogranin A as an endocrine marker, and glucagon and somatostatin to label α- and δ-cells, respectively. These markers were selected to assess the integrity of multiple endocrine cell populations and to determine whether prior DTZ staining interfered with downstream immunostaining or antigen detection. DTZ-treated grafts showed clear and specific labeling for all markers examined, with no apparent loss of signal or alteration in cellular distribution (Fig. 3D). Together, these results show that DTZ staining facilitates recovery of subcutaneously transplanted islet grafts without adversely affecting subsequent immunohistochemical assessment of pancreatic endocrine cell composition.

## DISCUSSION

By replacing endogenous insulin production, islet transplantation has the potential to eliminate the need for lifelong insulin therapy. However, despite advances in islet isolation and transplantation protocols, the long-term success of this therapy remains limited by issues such as graft loss, immune rejection, and insufficient methods for non-invasive graft tracking (11,29). Developing techniques that support better monitoring and recovery of islet grafts could accelerate studies aimed at optimizing cell-based therapies (30,31).

A central factor influencing graft outcomes is the choice of transplantation site (32–34). While the hepatic portal vein remains the clinical standard, this transplantation site poses challenges including early inflammatory responses, limited access for graft imaging, and poor long-term survival. In contrast, alternative sites such as the kidney capsule and intramuscular tissue are frequently used in preclinical models but present limitations in scalability and retrieval (35). The subcutaneous space offers a promising alternative due to its ease of access, minimal invasiveness, and compatibility with emerging clinical approaches. However, a key drawback has been the difficulty in reliably locating and retrieving engrafted islets, which our DTZ-based approach aims to directly address.

Beyond locating the ideal site for islet transplantation, improving islet graft survival is critical to advancing therapeutic efficacy (32,34,36). Immune rejection is one of the primary obstacles of islet graft survival, prompting strategies such as systemic immunosuppression or immune-isolating encapsulation to reduce immunogenicity (37–39). These approaches often require longitudinal assessment of graft function and persistence, which is difficult to achieve without non-invasive tracking tools. The ability to localize grafts using DTZ may provide further assessments of immune responses and rejection dynamics in preclinical studies.

Similarly, strategies to improve the local transplant environment, such as promoting vascularization or reducing inflammation, have shown promise in improving islet engraftment and function (11,40). However, evaluating these interventions often requires recovery and analysis of the transplanted islets, which has been challenging in sites like the subcutaneous space. The DTZ method proposed here helps streamline this process by enabling reliable identification and retrieval of grafts and supports ongoing efforts to improve the viability and scalability of islet cell therapy.

## CONCLUSIONS

In this study, we demonstrate that DTZ can be successfully applied post-transplantation to identify islet grafts in the subcutaneous space. DTZ produced clear and selective red staining of insulin-producing β-cells in both SC-derived and primary human islets grafts. These findings establish DTZ as a simple and cost-effective tool that helps facilitate graft monitoring and recovery. We further demonstrate the utility of the BioRep ICC automated image-based islet counter to quantify SC-islets before transplantation into mice.

## ACKNOWLEDGEMENTS

This work was supported by the NIH (R01DK114233, UG3DK142188), Breakthrough T1D (3-SRA-2023-1295-S-B), the Edward J. Mallinckrodt Foundation, the Anita Palmer Corbin Trust, and Washington University to J.R.M. M.I. was supported by the Rita Levi-Montalcini Postdoctoral Fellowship in Regenerative Medicine and the NIH (T32DK007120). M.M.M. was supported by the Cellular and Molecular Biology Training grant (T32GM139774). Further support was provided by the Washington University Diabetes Research Center (P30DK020579). We would like to thank Erika Brown (Washington University) for her support in graphic design on the manuscript.

## AUTHOR CONTRIBUTIONS

J.L., M.I., M.M.M., and J.R.M. designed the project and experiments. J.L., M.I., and J.R.M wrote the manuscript. All authors revised and approved of the manuscript.

## COMPETING INTERESTS

M.I. has stock in Vertex Pharmaceuticals. J.R.M. is an inventor of licensed patents and patent applications, is the founder of IsletForge Innovations, Inc., and has stock in Sana Biotechnology. The remaining authors declare no competing interests.

## LIST OF ABBREVIATIONS

T1D: Type 1 Diabetes
SC-islets: Stem cell-derived islets
Kidney Capsule: renal subcapsular space
IEQ: Islet equivalents
BioRep ICC: BioRep Islet Cell Counter
DTZ: Dithizone
IACUC: Insitutional Animal Care and Use Committee
KRB: Kreb’s Ringer Bicarbonate Buffer
GTT: Glucose tolerance test
GSIS: Glucose stimulated Insulin Secretion
IHC: Immunohistochemistry

